# Principles of RNA recruitment to viral ribonucleoprotein condensates in a segmented dsRNA virus

**DOI:** 10.1101/2021.03.22.435476

**Authors:** Sebastian Strauss, Alexander Borodavka, Guido Papa, Daniel Desiró, Florian Schueder, Ralf Jungmann

**Author notes:** Correspondence should be addressed to A.B. and R.J. These authors contributed equally to this work (Sebastian Strauss, Alexander Borodavka). Medical Research Council Laboratory of Molecular Biology (MRC LMB), Cambridge Biomedical Campus, Francis Crick Avenue, Cambridge CB2 0QH, UK.

## Abstract

Rotaviruses transcribe eleven distinct protein-coding RNAs that must be stoichiometrically co-packaged prior to their replication to make an infectious virion. During infection, rotavirus transcripts accumulate in cytoplasmic ribonucleoprotein (RNP) condensates, termed viroplasms. Understanding the mechanisms of viroplasm assembly and RNA enrichment within is crucial to gaining greater insight into their function and stoichiometric assortment of individual transcripts. We analysed the subcellular distribution of individual RV transcripts and viroplasm transcriptome by combining multiplexed DNA-barcoded single-molecule RNA FISH of infected cells. Using DNA-PAINT microscopy, we provide evidence of the early onset of viral transcript oligomerisation that occurs prior to the formation of viroplasms. We demonstrate that viral sequences lacking the conserved terminal regions fail to undergo enrichment in rotavirus RNP condensates. We show that individual viral transcripts exhibit variable propensities to partition into viroplasms, irrespective of their absolute numbers in cells, suggesting a selective RNA enrichment mechanism distinct from other known cellular RNP granules. We suggest that rotavirus replication factories represent unique RNP condensates enriched in eleven types of cognate transcripts that may facilitate the assembly of a multi-segmented RNA genome.

## Introduction

RNA genome segmentation poses challenges for assembly and genome packaging of viruses, including rotaviruses, a large group of human and animal pathogens. The rotavirus (RV) genome is enclosed in a protein shell, inside which eleven double-stranded (ds)RNAs, also known as genomic segments, iteratively undergo rounds of transcription. Consequently, multiple copies of distinct RV transcripts accumulate in the cytoplasm of infected cells^1–6^. It remains a long-standing mystery how RVs robustly select and co-package eleven non-identical RNAs despite the non-stoichiometric transcript accumulation in cells^7^. Within 2 hours of the onset of transcription, viral RNA-binding proteins NSP2 and NSP5^8–10^ begin to concentrate within membraneless cytoplasmic replication factories, also known as viroplasms^11,12^. Experimental evidence suggest that viroplasms may provide a selective environment preventing siRNA-targeted degradation of the RV transcripts^13^ that serve as templates for the synthesis of the viral dsRNA genome. Previous attempts to investigate the ultrastructure of viroplasms have not succeeded in revealing the identities and stoichiometry of individual transcripts therein^12,14–16^, and it thus remains unclear whether these granules contain all eleven non-identical RNA species, and if so, how these organelles maintain their unique RNA composition.

Recently, we have discovered that early replication stage (2-6 hours post infection) viroplasms represent liquid condensates that are formed via phase-separation of the RV RNA chaperone NSP2 and a non-structural phosphoprotein NSP5^17^. These condensates could be rapidly and reversibly dissolved by treating RV-infected cells with small aliphatic diols^17^. A fraction of the RV transcripts was released from viroplasms when they were briefly treated with 4.5% propylene glycol or 4% 1,6-hexanediol, resulting in the reversible recruitment of the RV transcripts into these condensates when these compounds were removed^17^. In principle, the formation of dynamic ribonucleoprotein condensates in RV-infected cells could facilitate selective enrichment of cognate viral transcripts required for a robust and stoichiometric RNA assembly and packaging. Despite the extensive evidence of the importance of viroplasms in RV replication^13,18–22^, the analysis of their molecular composition have been confounded by both their dynamic and liquid-like nature that precluded successful isolation from the RV-infected cells. Thus, the exact RNA composition of these assemblies has remained enigmatic.

To unravel the principles of viral transcript assembly and partitioning into the rotavirus RNP granules, we have visualised the RV transcriptome using a DNA barcode-based multiplexing approach^23^, combined with single-molecule RNA Fluorescence In Situ Hybridisation (smFISH). We have shown that during early infection, RV transcripts are detected as diffusely distributed species, albeit with different efficiencies, suggesting their non-stoichiometric production. We used RNA-Seq to quantify the transcriptome of RV-infected cells at 6 hours post infection, revealing the extreme abundance of viral transcripts constituting approximately 17% of all detected protein-coding transcripts. Multiplexed smFISH quantification of single infected cells broadly agrees with the RNA-Seq data, confirming their non-stoichiometric accumulation in cells. Using super-resolution DNA-PAINT^24^ and molecular counting qPAINT^25^ approaches, we have demonstrated that single RV transcripts first aggregate to form cytoplasmic RNA clusters that precede the formation of detectable viral replication factories. Silencing of the viroplasm-forming RNA chaperone NSP2 decreased the propensities of viral transcripts to associate. Together with previous *in vitro* studies^26,27^, these observations present new supporting evidence of NSP2 acting as an RNA chaperone required for rotavirus transcript assembly. Multiplexed smFISH imaging of individual viral ribonucleoprotein granules has revealed that all eleven types of transcripts are highly enriched irrespective of their apparent cytoplasmic stoichiometry. To further understand the selectivity of the observed transcript enrichment in viroplasms, we generated a recombinant rotavirus harbouring an EGFP-coding insertion in a viral gene segment, to investigate the effect of non-viral heterologous sequences on the RNA localisation. These EGFP-coding transcripts were enriched in viroplasms, while similar EGFP-coding mRNAs lacking segment-specific conserved terminal regions did not partition into these membraneless organelles. These results reveal key differences in the mechanisms of RNA partitioning that underlie the assembly of other membraneless RNA-rich granules, e.g., stress granules and P-bodies^28–30^, and viroplasms. We suggest that such a mechanism of RNA partitioning into complex biomolecular condensates in a multi-segmented RNA virus would facilitate the enrichment of non-identical RV transcripts required for their RNA-chaperone assisted stoichiometric assembly within these ribonucleoprotein condensates.

## Results

### NSP5-EGFP-tagged Viral Condensates Retain Viral Transcripts

In order to monitor rotavirus transcription during the course of infection, we took advantage of the MA104 cell line stably expressing an EGFP-tagged NSP5 that readily partitions into the viroplasmic condensates^18,31,32^. At a multiplicity of infection (MOI) of 10, NSP5-EGFP-tagged granules were readily detected as soon as 2–3 hours post infection (hpi) (**Fig. 1**). Recently, we have shown that during these early infection stages, such NSP5-rich granules exhibit liquid-like behaviour, representing dynamic NSP5:NSP2 condensates^32^. To confirm that these EGFP-NSP5-tagged condensates represent *bona fide* viroplasms that accumulate viral RNA, we examined the RV transcript accumulation by Fluorescence in Situ Hybridisation (FISH). We used a pooled set of probes consisting of 3 oligonucleotides targeting protein-coding sequences of each segment of the Rotavirus A genome (G6P6[1] strain RF, further details of probes and fluorophores – see **Supplementary Table S1** and **Supplementary file 1).** Multiple RNA-rich foci were readily detected in cells (**Fig. 1**) as early as 3 hpi, with high degree of colocalisation with NSP5-EGFP signal (**Fig. 1c**). Given that each RV transcript is targeted by 3 transcript-specific probes, and the observed point sources could only be detected after 2-3 hpi, these signals are unlikely to originate from the hybridisation events to single transcripts^33,34^. All detectable newly-formed NSP5-EGFP-tagged condensates retained RV transcripts (**Fig. 1c**), further suggesting that these condensates represent sites of RV replication, consisted with their proposed role in supporting viral replication^12^. During the course of infection, a fraction of the viral transcriptome was also detected outside the NSP5-EGFP-tagged condensates, resulting in lower degree of the RV RNA/EGFP-NSP5 colocalisation (**Fig. 1c**). However, no RV-specific RNA signal was detected in the RV-infected cells up to 3 hpi (**Fig. 1**), confirming the specificity of the designed 3x probes towards the RV transcripts. Given that EGFP-tagged viroplasms accumulate large amounts of viral RNA-binding proteins^35,36^ known to promiscuously bind non-viral RNAs^27,35^, we then explored whether other non-viral, highly expressed cytoplasmically localised transcripts would undergo enrichment in these granules. We performed smFISH to visualise GAPDH transcripts (**Supplementary Fig. S1**). The apparent intensity distribution for GAPDH signals was unimodal, as expected for single non-interacting transcripts. A number of cells also contained rotavirus RNA-specific foci that co-localised with EGFP-tagged viroplasms at 4 hpi (**Fig. 1** and **Supplementary Fig. S1**). Conversely, the high-copy number GAPDH transcripts did not undergo enrichment in viroplasms, suggesting an RNA selection mechanism that determines transcript partitioning into these viral organelles.

**Figure 1.**
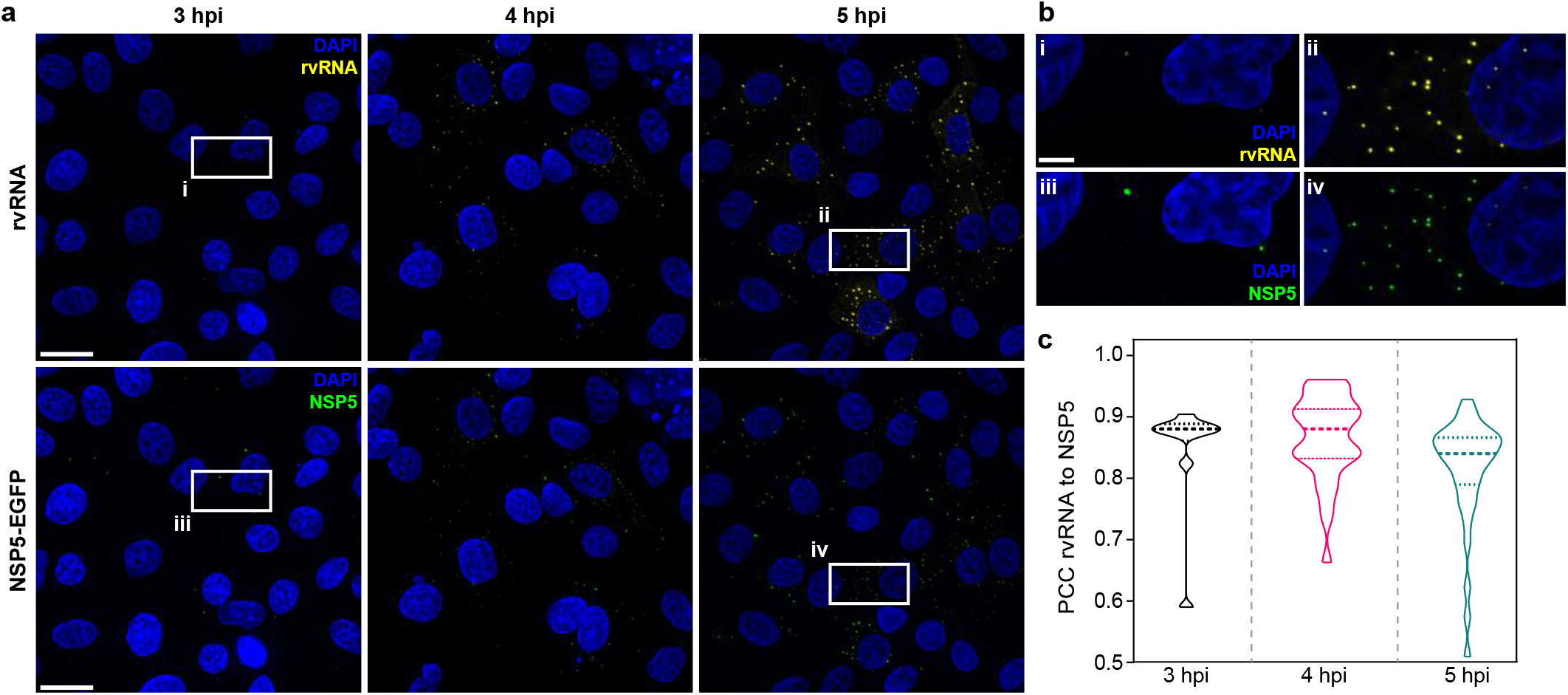
RNA Accumulation During Rotavirus Infection. (**a**) RV RNA accumulation in viral factories during early infection. Top: RV transcripts (yellow) visualized by smFISH using a combined set of 3-5 segment-specific probes targeting each genomic transcript. Bottom: NSP5-EGFP-tagged viral factories (green). Images represent maximum intensity Z-projections acquired using identical settings and brightness levels for both channels. DAPI-stained nuclei are shown in blue. Scale bars, 25 μm. (**b**) RV transcripts and NSP5-EGFP-specific signals shown in the white box in panel **a**. Note the low intensity of the RNA and EGFP signals during early infection stages that increase by 5 hpi. Scale bar, 5 μm. (**c**) Pearson’s correlation coefficients (PCCs) of colocalising NSP5-EGFP tagged viral factories and RV transcripts, as shown above in panel (**b**). Dashed and dotted lines represent median and quartile values of PCCs, respectively. Each data point represents the PCC value calculated for a single cell. N = 10 (3 hpi), N = 31 (4 hpi), N = 38 (5 hpi).

To discern gene-specific viral transcripts, we designed two distinct sets of single-molecule (sm)FISH probes, each targeting the coding regions of the RV gene segments (Seg) Seg3 and Seg4 transcripts, respectively. Each pool consisting of 48 spectrally distinct probes (**Materials and Methods**) generates high-intensity spots upon hybridisation to an individual transcript. At 2 hpi, both Seg3 and Seg4 transcripts were readily detectable as point sources that were sufficiently far apart to be resolved as individual points (**Fig. 2**). The observed uniformity of intensities of point sources (**Supplementary Fig. S2**) was comparable to the GAPDH RNA signal distribution visualised using a set of smFISH probes labelled with an identical fluorophore under the same imaging conditions (**Materials and Methods**), further confirming that these objects represented single RV transcripts. Both Seg3 and Seg4 transcripts were equally abundant and randomly distributed in the cytoplasm of infected cells without a discernible pattern. Such random point distribution further suggested a lack of directional transport of RNAs^33^ away from the transcribing viral particles.

**Figure 2.**
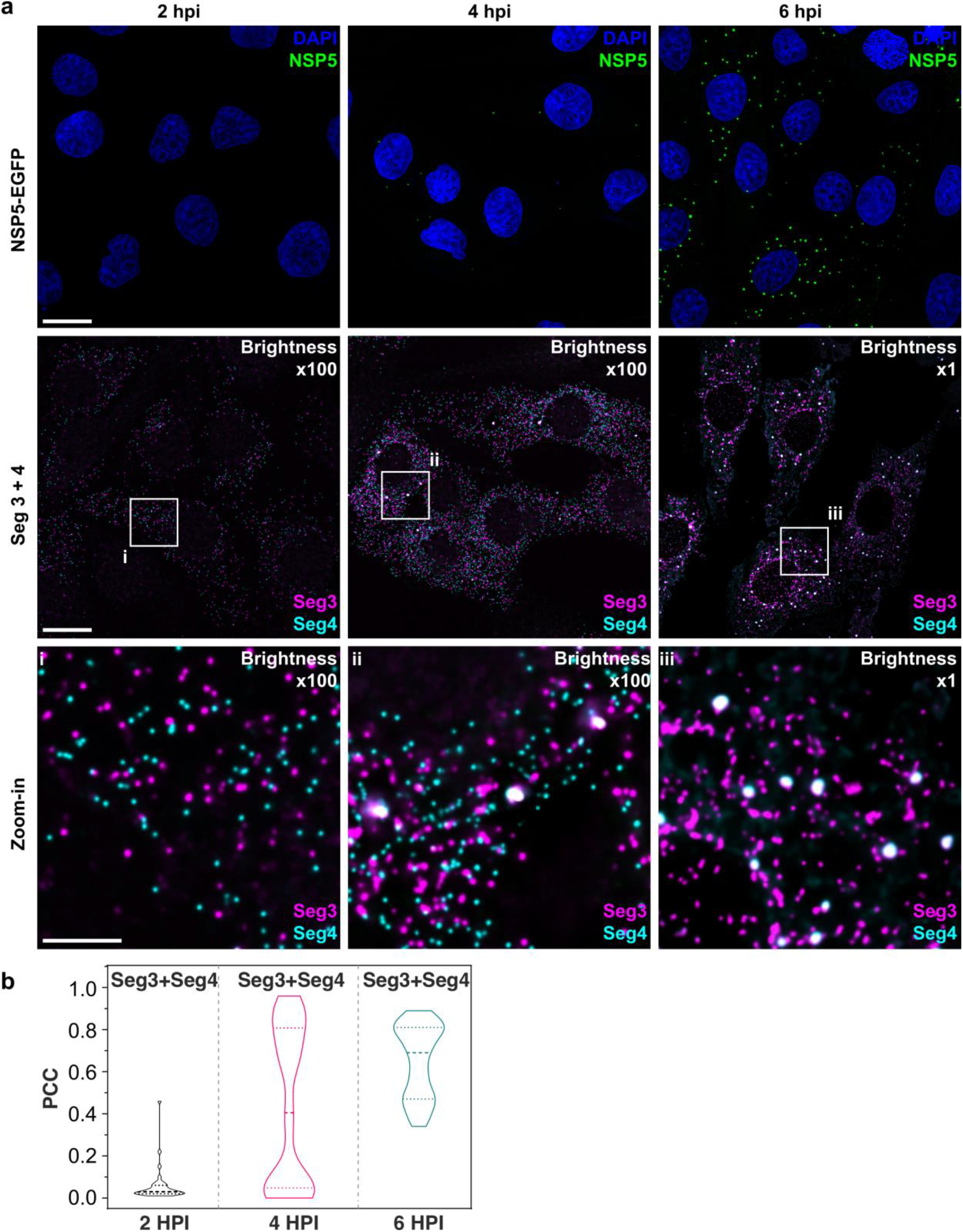
Segment-specific Transcript Accumulation During Rotavirus Infection. (**a**) smFISH of RV transcripts at early infection time points (2–6 hpi) using segment-specific probes. Two sets of 48 gene-specific probes were designed for each Seg3 and Seg4 RNA transcripts. Top: NSP5-EGFP-tagged viral factories (green), nuclei (blue). Middle: Seg3 (magenta) and Seg4 (cyan) RNA signals in the ROIs shown above, with colocalising Seg3 and Seg4 RNAs (white). As the amount of each transcript increases during the course of infection, brightness settings were adjusted accordingly to reveal single transcripts (Brightness x1000 in the panels above and 1x in the panel (**b**) for comparison). Scale bars, 25 μm (**a**), 5 μm (**b**). (**c**) Colocalisation of Seg3 and Seg4 transcripts. Pearson’s correlation coefficients (PCCs) of colocalising Seg3 and Seg4 transcript signals. Dashed and dotted lines represent median and quartile values of PCCs, respectively. Each data point represents the PCC value calculated for a single cell. N = 37 (2 hpi), N = 62 (4 hpi), N = 51 (6 hpi). Statistical analysis of data was performed using Kolmogorov-Smirnov test. At the 0.01 level, the observed distributions are significantly different.

### Rotavirus Transcript Association Requires RNA Chaperone NSP2

The overall cytoplasmic density of transcripts increased over time, reflecting ongoing viral transcription (**Fig. 2** and **Supplementary Fig. S3**). A hallmark of the rotavirus replication cycle is an exponential increase in the amount of RNA produced after 4-6 hpi^7,37^, emanating from the second round of transcription by the newly assembled particles. We therefore initially focused on analysing the intracellular distribution of viral transcripts between 2-3 hours post infection. Despite the apparently equal ratio of Seg3 and Seg4 transcripts produced between 2-4 hpi (**Supplementary Figs. S2** and **S3**), at 6 hpi the amount of Seg3 was significantly higher than that of Seg4 RNA (**Supplementary Fig. S3**), suggesting that individual viral transcripts have different half-lives in the cytoplasm. After 3 hpi, multiple Seg3 and Seg4 transcripts co-localised, resulting in much higher intensity signals compared to single transcripts (**Fig. 2** and **Supplementary Fig. S2** and **S3**). The number of high-intensity RNA foci further increased between 3-6 hpi, manifesting in a higher density of co-localising Seg3 and Seg4 transcripts (**Fig. 2**). We hypothesised that the observed RNA aggregation depends on the production of viral RNA-binding proteins^30^. A large number of Seg3 and Seg4 transcript-containing clusters co-localised with EGFP-tagged condensates (**Supplementary Fig. S4**), highly enriched in a multivalent RNA chaperone NSP2^6,12,38^. Previously, we have shown that NSP2 may facilitate inter-molecular RNA association *in vitro*^26,27^. To further investigate the role of NSP2 in the observed RNA aggregation, we analysed two cell lines, each of which was expressing a short-hairpin RNA (shRNA) targeting either the NSP2 gene, or a scrambled control RNA. At 4 hpi, shRNA-mediated NSP2 knockdown resulted in an overall reduction of signal intensities for both Seg3 and Seg4 RNAs (**Supplementary Fig. S5**). Importantly, NSP2 knockdown (**Supplementary Fig. S6)** disrupted the apparent aggregation of Seg3 and Seg4 transcripts that no longer formed high-intensity RNA foci (**Supplementary Fig. S5**). Together, these data indicate that Seg3 and Seg4 RNA clustering requires expression of the viral RNA chaperone NSP2, further suggesting the role of NSP2 in mediating the formation of higher order viral transcript assemblies. To be able to directly visualise transcript oligomerisation in cells, we carried out super-resolution imaging of individual RNAs, and their oligomers using smFISH combined with DNA-based point accumulation for imaging in nanoscale topography (DNA-PAINT) approach^24,39^. We carried out a quantitative qPAINT analysis^25^ (**Fig. 3**) of Seg3 transcripts to assess the approximate number of its RNA-binding sites at 2 hpi (early infection stage) when the density of transcripts is low. The qPAINT analysis (Materials and Methods and **Supplementary Table S2**), revealed an apparent *k_on_* of 10^7^ (Ms)^−1^ that corresponds to approximately ten FISH probes per transcript^25^, consistent with these structures being single transcripts. Between 4-6 hpi, a fraction of Seg3 transcripts underwent assembly, yielding larger RNA clusters (**Fig. 3**) that contained approximately 20-50 transcripts. Together, these results indicate that Seg3 transcript clustering is concomitant with the observed viral RNA aggregation during infection, and this process requires expression of the viral RNA chaperone NSP2.

**Figure 3.**
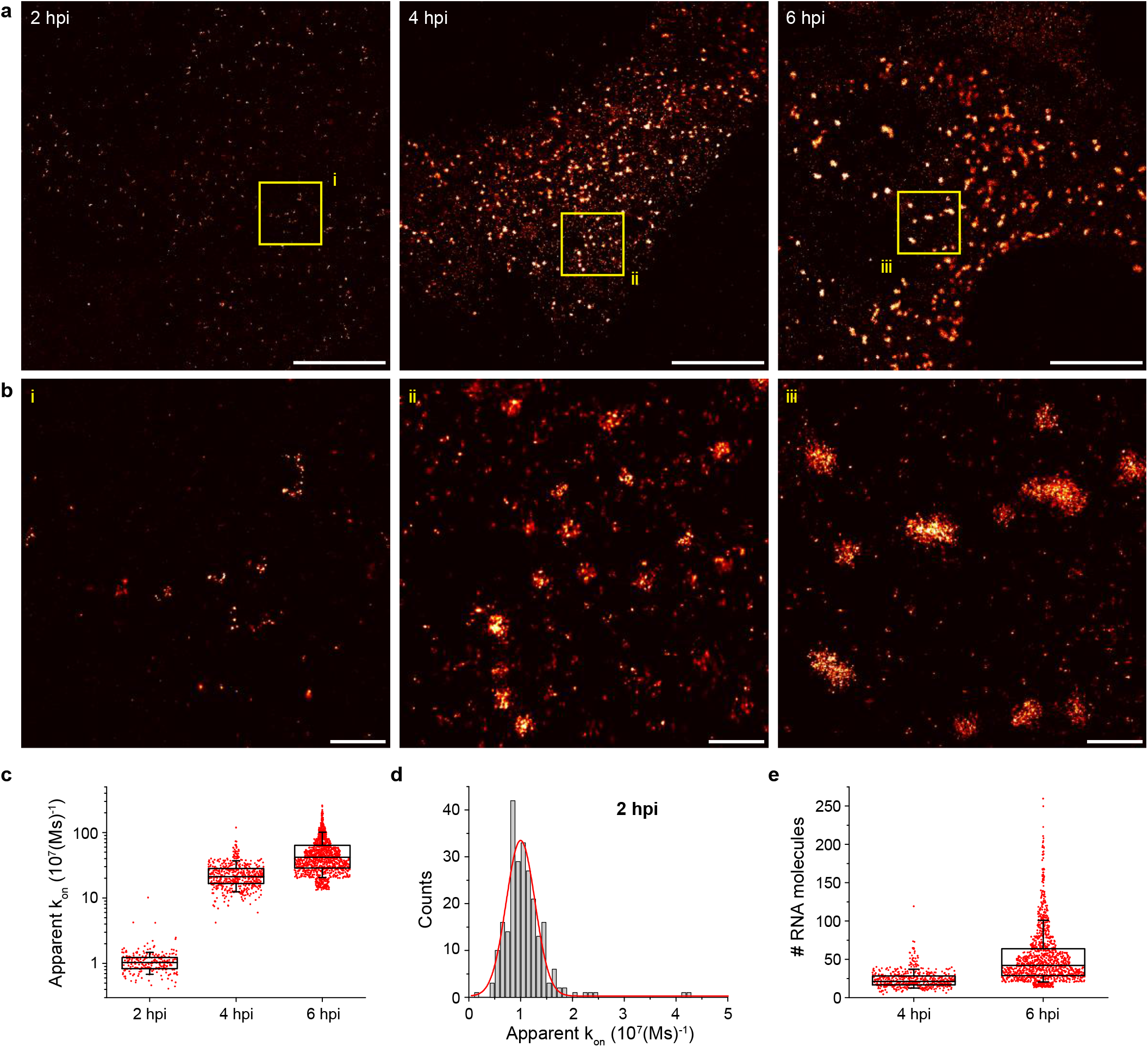
DNA-PAINT Analysis of the RV RNA Oligomerisation. (**a**) DNA-PAINT super-resolution micrograph of Seg3 transcripts at 2, 4 and 6 hpi. (**b**) Zoomed-in regions of Seg3 RNA structures highlighted in (**a**). (**c**) Quantitative PAINT (qPAINT) analysis of the RNA structures shown above. *k*_*on*_ values were calculated for each selected structure assuming *k*_*on*_ = (*τ*_*D*_ × *c*)^−1^, where *c* is the imager strand concentration and *τ*_*D*_ the dark time between binding events (c_imager_ = 1 nM at 2 hpi; 125 pM at 4 hpi and 100 pM at 6 hpi). Each point represents the apparent *k_on_* value for an RNA structure that is directly proportional to a relative number of the FISH probe binding sites, and thus reflects the number of transcripts within. (**d**) Unimodal distribution of *k_on_* values calculated for single Seg3 transcripts detected at 2 hpi. The resulting average k_on_ value calculated from a Gaussian fit is 10^−5^ (Ms)^−1^, corresponding to a single Seg3 transcript. (**e**) Estimated numbers of Seg3 transcripts (mean±SD) at 4 hpi and 6 hpi are 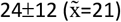 and 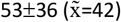, respectively. N = 245 (2 hpi), N = 515 (4 hpi), N = 1181 (6 hpi). Scale bars, 5 μm (**a**), 500 nm (**b**).

### Single-cell Rotavirus Transcriptome Analysis Using Universal DNA Exchange and smFISH (UDEx-FISH)

To be able to visualise the RV transcriptome in single cells, we employed Universal DNA Exchange approach^23^ combined with smFISH (termed ‘UDEx-FISH’) to sequentially image individual transcripts (Seg1-Seg11). Eleven RNA targets of interest were first pre-hybridised with each set of probes containing transcript-specific sequences that stably hybridise with the RNA, followed by a shorter a ‘handle’ sequence that binds fluorescently labeled complementary DNA probes (‘Imager’, **Fig. 4a**). This approach allows installation of unique DNA barcodes onto individual transcripts, thus enabling multiplexed single-molecule imaging of targets irrespective of their molecular density unlike alternative combinatorial labelling schemes, e.g., MERFISH^40^.

**Figure 4.**
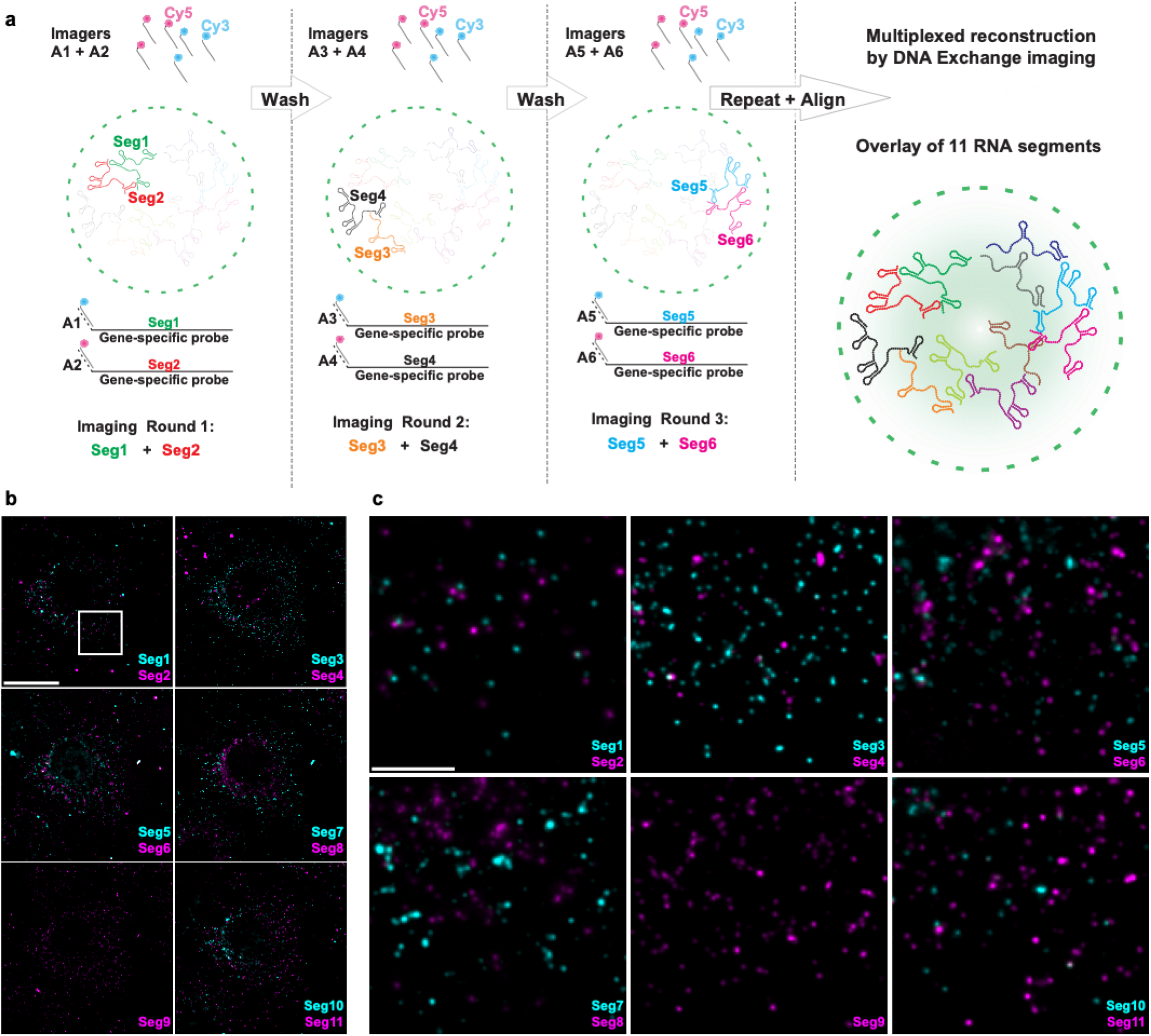
Universal DNA Exchange-smFISH Approach for Single-Cell Imaging of the RV Transcriptome. (**a**) Schematics of the Universal DNA Exchange-smFISH approach (UDEx-FISH). FISH probes targeting different RNA segments are extended with 12 nucleotide-long orthogonal DNA sequences A_1_-A_n_ (DNA ‘handles’). Cy3 and Cy5-dye-modified DNA sequences (A1–An, ‘Imagers’) complementary to their respective handles can rapidly (<1 min) and stably hybridize with their segment-specific FISH probes RNA. Non-hybridized imagers are washed away and the two RNA targets, as well as NSP5-EGFP tagged viroplasms and nuclei are imaged in each imaging round. Afterwards, the two imager DNA strands are removed under gentle denaturing conditions (see Materials and Methods for details), allowing the next imaging round to take place. The imaging procedure is repeated until all targets of interest are recorded without affecting the quality of the fixed sample or loss of the RNA signal intensity due to photobleaching. (**b**) RV Transcriptome during early infection stage (2 hpi). Images were taken with same acquisition parameters for each imaging round. Scale bar, 20 μm. (**c**) Zoomed-in images of the highlighted areas shown in (**b**). Scale bar, 5 μm.

An equimolar mixture of transcript-specific pools of UDEx-FISH probes was hybridised with cells at different points of the viral infection. RV transcriptome was imaged via successive rounds of a brief incubation with target-specific fluorescently labelled DNA imagers, followed by their removal during a formamide-containing wash step (**Fig. 4b**). To ascertain that during wash steps only DNA imagers were removed without the loss of transcript-specific smFISH probes, we carried out 5 iterative washes/imager applications. The relative fluorescence intensities of transcript-specific imagers remained unchanged (**Supplementary Fig. S7**) after 5 cycles of washes. More importantly, no residual signal was recorded after each individual wash step (**Supplementary Fig. S7**), and no transcript signal loss was observed due to bleaching. Finally, multiple rounds of washes did not alter the distribution of high-intensity RNA foci, nor had any apparent impact on the distribution or the morphology of RNA clusters (**Supplementary Fig. S7**), or EGFP-tagged condensates, confirming that the chosen approach and the designed probes were highly suitable for multiplexed characterisation of the RV transcriptome in single cells. At 2 hpi, multiple copies of each transcript Seg1-Seg11 were readily detected, prior to the formation of EGFP-tagged viral replicative condensates (**Fig. 4b**). Assuming similar rates of transcription for each individual genomic segment^1,7^, transcription of longer segments is expected to yield fewer copies of longer Seg1-Seg4 transcripts (3.4-2.6 kb). Surprisingly, Seg3 transcript was more abundant than similarly sized Seg2 or Seg4 transcripts, suggesting that individual viral transcripts may have different stabilities resulting in distinct kinetics of transcript accumulation during infection.

### Transcript Partitioning into Viral Condensates is Influenced by Their Untranslated Terminal Regions

We then examined the intracellular distribution and assembly of the remaining RV transcriptome during the onset of formation of detectable NSP5-rich condensates. At 4 hpi, NSP5-EGFP tagged condensates contained all eleven types of RV transcripts (**Figs. 5a**-**b**), confirming that the observed cytoplasmic inclusions represent genuine viral ribonucleoprotein condensates. RNA-Seq quantification of transcripts revealed that as early as 4-6 hpi, the RV transcriptome represented approximately 17% of all mapped protein-coding transcripts in cells (**Fig. 5c and Supplementary file 2**). Overall, Seg11, Seg6, Seg10 and Seg7 transcripts were the most abundant viral RNA species quantified by RNA-Seq (**Fig. 5c**). While single-cell smFISH RNA quantification was broadly in agreement with these data, we have noted that the smFISH detection efficiency of Seg11 transcripts was low, likely due to the smallest size of its coding region (< 0.6 kb), as well as a potentially poor accessibility of the RNA target^26,41^. In contrast, the largest (3.4 kb) Seg1 transcripts were the least abundant viral RNA species quantified by both RNA-Seq and smFISH (**Figs. 5c-d**). Interestingly, despite the lowest abundance in the RV transcriptome, Seg1 RNAs were the most enriched species in the viral RNP condensates (**Fig. 5e**). Similarly, despite the smallest (<0.8 kb) sizes, thus lower detection efficiencies, Seg10 and Seg11 transcripts were also efficient in undergoing enrichment in viroplasms (**Fig. 5e**), revealing that RNA partitioning into these NSP2/NSP5-rich condensates does not simply reflect their GC content^29^, cytoplasmic abundance, or size^28,42,43^, and it is likely to be determined by other factors, e.g., segment-specific untranslated regions (UTRs). To further explore this possibility, we examined whether NSP5-EGFP transcripts containing the coding region of Seg11 RNA (NSP5) produced in a stable NSP5-EGFP MA104 cell line would undergo enrichment in viroplasms during RV infection (**Fig. 6a**). As expected, all cells were positive for EGFP-coding NSP5-EGFP chimeric transcripts that were diffusely distributed in the cytoplasm of RV-infected cells at 8 hpi. Abundant NSP5-EGFP-tagged viroplasmic RNP condensates confirmed that NSP5-EGFP transcripts retained their coding capacity (**Fig. 6b,** top panel). However, despite the presence of the coding NSP5 sequence, these polyadenylated transcripts devoid of segment-specific UTRs did not undergo enrichment in viroplasms (**Fig. 6b**). To investigate whether the inclusion of a non-viral, heterologous EGFP sequence might affect the enrichment of a cognate viral RNA sequence, we also visualised an EGFP-coding sequence inserted into a gene segment 7 that contained intact segment-specific UTRs (**Fig. 6a**). These viral EGFP-coding transcripts were highly enriched in the NSP5-EGFP-tagged viroplasmic condensates that formed during infection with a recombinant NSP3-2A-EGFP virus (**Fig. 6,** bottom panel). Collectively, these results strongly suggest that segment-specific UTRs determine the capacity of the RV transcripts to undergo enrichment in viral condensates.

**Figure 5.**
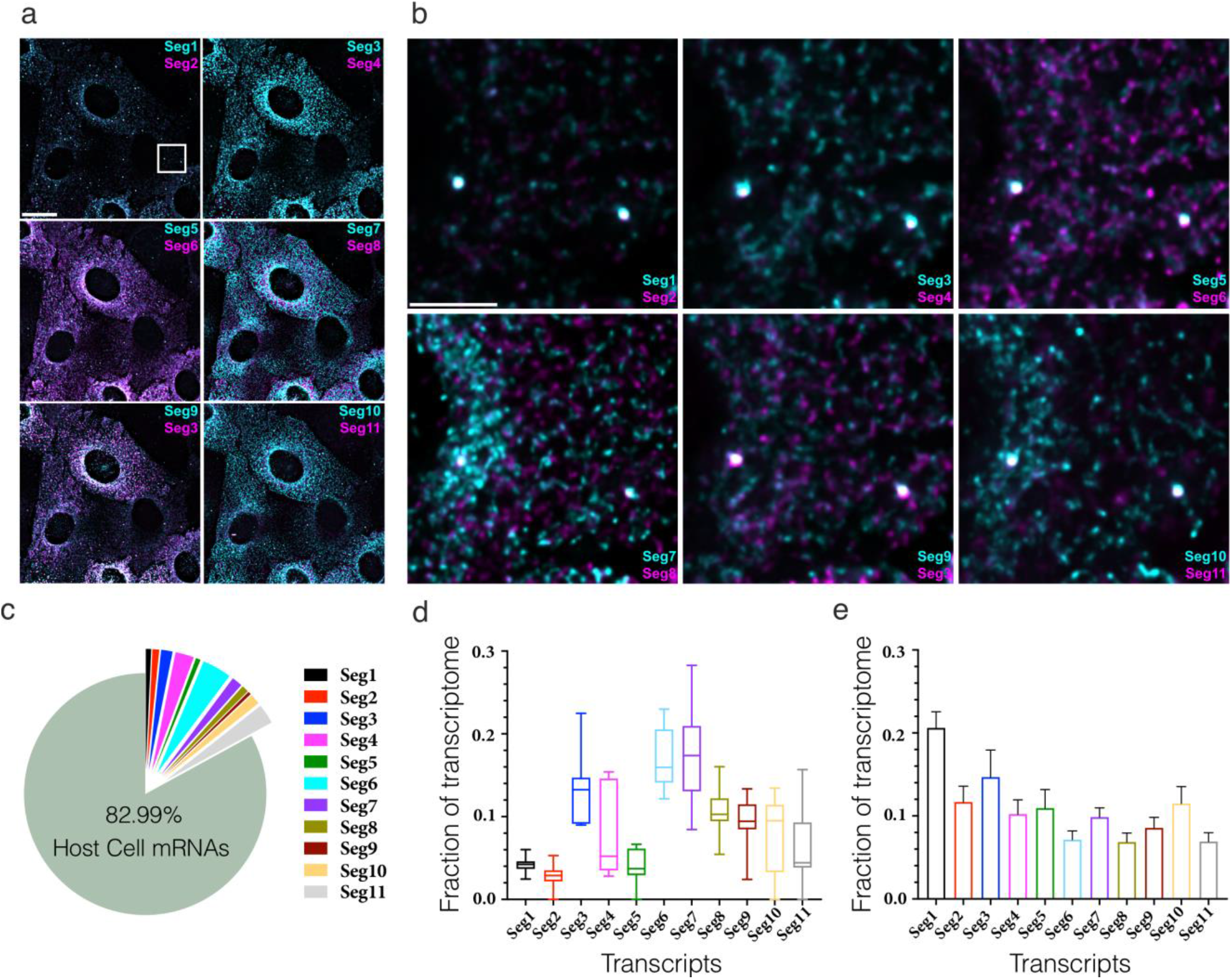
Single-cell Analysis of the RV Transcriptome. (**a**) *In situ* analysis of the RV transcriptome at 6 hpi. Scale bar, 20 μm. (**b**) Zoomed-in images of the highlighted regions containing viroplasms and 11 distinct RNA targets. Scale bar: 5 μm. (**c**) The proportion of RNA-seq reads assigned to the annotated protein-coding transcriptome of MA104 cells infected with RVA strain RF at 6 hpi. The RVA transcriptome represented ~17% of total reads mapped by Salmon (*See Materials and Methods*). (**d**) Single-cell analysis of the relative fractions of each segment-specific transcript (N = 10 cells) estimated from the UDEx-FISH images data shown in panel (a). (**e**) Relative fractions (Mean ± StDev) of individual viral transcripts detected in individual NSP5-EGFP-rich granules detected in 10 cells.

**Figure 6.**
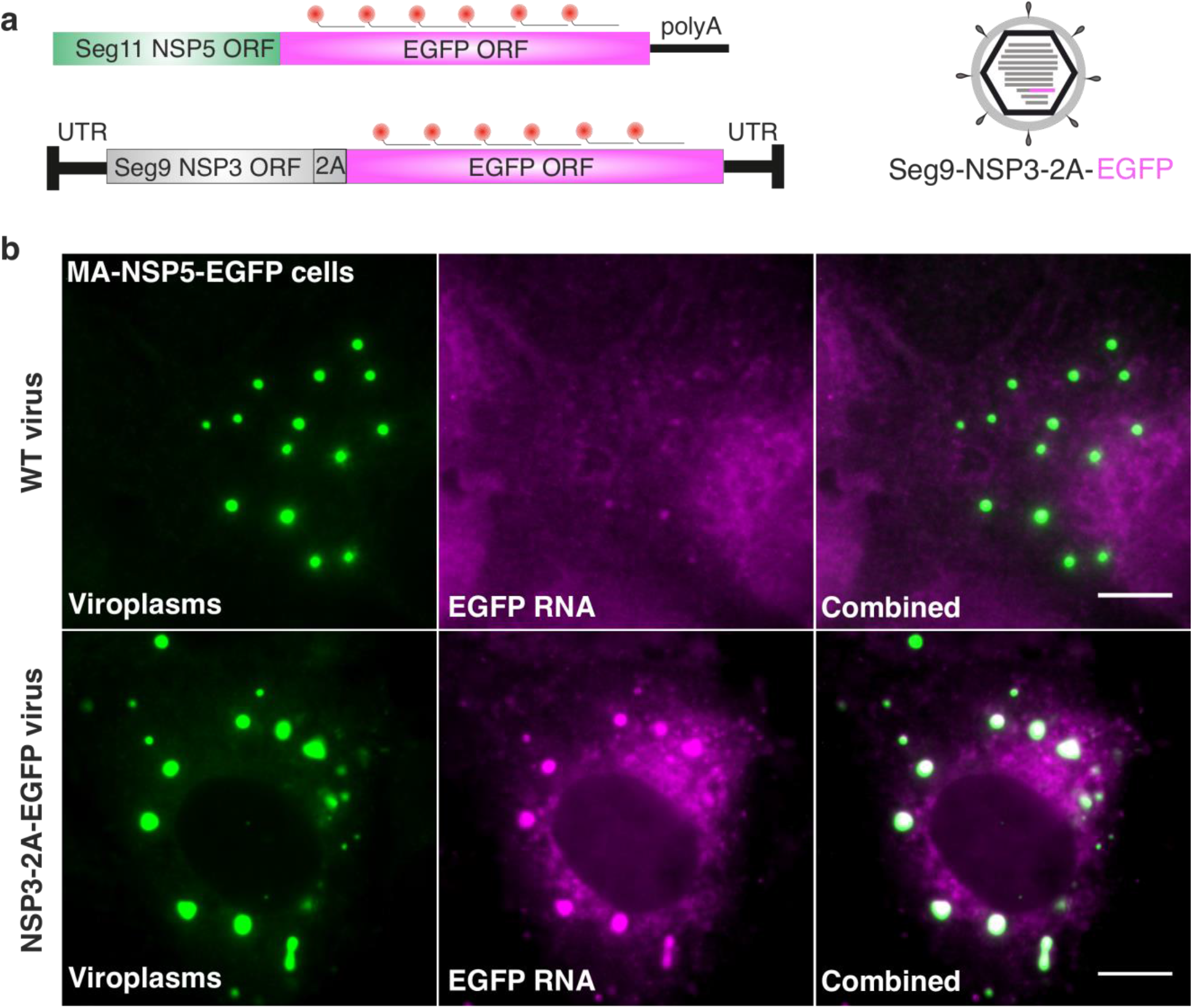
EGFP-coding sequences lacking segment-specific UTRs do not localize to viroplasms. (**a**) Experimental design of smFISH probes to distinguish chimeric rotavirus RNA fused to EGFP-coding sequences transcribed in the nucleus (polyadenylated NSP5-EGFP coding mRNA), and transcribed by the recombinant virus NSP3-2A-EGFP (Seg9 RNA coding NSP3-2A-EGFP, containing intact segment-specific UTRs). (**b**) **Top panel** - MA-NSP5-EGFP cells infected with wild type rotavirus, fixed at 8 hpi. NSP5-EGFP-marked viroplasms (green) do not accumulate polyadenylated NSP5-EGFP-coding transcripts constitutively expressed in this cell line (EGFP RNA FISH, magenta). **Bottom panel** - MA-NSP5-EGFP cells infected with the recombinant rotavirus NSP3-2A-EGFP. An identical EGFP-coding sequence inserted into gene segment 9 of rotavirus results in enrichment of EGFP-coding Seg9 transcript in viroplasms at 8 hpi. Maximum intensity Z-projections of both channels are shown. Scale bar, 10 μm.

## Discussion

Rotavirus replication factories, or viroplasms, have long been known as membrane-less organelles that emerge in the cytoplasm shortly after infection^12^. Our recent findings have revealed that these viral organelles are formed via liquid-liquid phase separation (LLPS) of the RV RNA-binding proteins NSP2 and NSP5^17^. Similar liquid-like membrane-less RNP condensates are ubiquitous in cells^42,44–47^; however, each specific type of an RNP condensate being unique in their protein and RNA composition that can be dynamically modulated by various processes. While some of the key protein constituents of such RNP condensates have been uncovered, their unique RNA composition often remains poorly characterised due to the challenges related to their isolation and purification. Here, we provide the first glimpse at the unique RNA composition of the rotavirus RNP condensates that support replication of an eleven-segmented RNA genome.

RNA-Seq and single-cell UDEx-FISH analyses have revealed that at 6 hpi rotavirus infection resulted in non-stoichiometric accumulation of distinct viral transcripts, in agreement with previous bulk kinetic studies of RV replication carried out during late infection^7,8,13^. Early infection (2 hpi) RV transcriptome analysis by UDEx-FISH shows that all eleven types of the RV transcripts were diffusely distributed in the cytoplasm of infected cells. At 6 hpi, Seg1 (VP1-coding) transcripts represented the smallest fraction of the RV transcriptome. While RNA-Seq quantification provided the relative ratios of viral and non-viral protein-coding transcripts, it failed to reveal specific RNA localisation and subcellular distribution in individual RV-infected cells. In contrast, our UDEx-FISH analysis appeared to undercount smaller RNA targets (Seg11 and Seg10), presumably due to a limited number of smFISH probes successfully hybridized to these transcripts. We therefore used both approaches to quantify the viral transcriptome in RV-infected cells. The initial transcription stage was followed by the apparent formation of higher order assemblies of the RV transcripts during a process that required the viral RNA chaperone NSP2. Our observations are fully consistent with previously reported inhibition of genome replication and virion assembly resulting from the loss of NSP2 expression^13^, providing new evidence for NSP2 as a multivalent RNA chaperone that links together populations of viral transcripts. In light of our recent findings^17^, these results reveal several parallels between viroplasmic condensates and other cytoplasmic RNP granules, including stress granules (SGs) and P-bodies. Both SGs and viroplasms represent liquid-like RNP assemblies that form from untranslating mRNAs^28,30,42^. Expression of SG-specific or viroplasm-specific multivalent RNA-binding proteins is essential to their formation. While many aspects of viroplasm formation mimic those seen in assembly of stress granules^28,30,42,48^, in contrast, RNA partitioning into viroplasms and the observed clustering of transcripts appears to be virus-specific, implying that both processes depend on cognate, RV-specific RNA-protein and RNA-RNA interactions. Unlike SGs, whose RNA composition is biased toward larger, AU-rich mRNAs^42^, our data reveal that viroplasmic RNA enrichment is likely to be determined by transcript-specific terminal sequences of each RV gene segment. It remains to be determined how the individual sequences within the transcript-specific UTRs of different length and composition determine RNA partitioning into these condensates. A plausible model for such enrichment is based on a high affinity, specific protein-RNA recognition, such as previously reported conserved interaction between the 3’-terminal sequence of each RV transcript and its RNA-dependent RNA polymerase (RdRP) exhibiting high nM affinity for NSP5^49–51^. We suggest that such unique RdRP binding sites located within the 3’ UTR of the viral transcripts would facilitate their selection and enrichment within the viral NSP5/NSP2-rich RNP condensates. Interestingly, a similar RdRP-facilitated viral transcript selection mechanism was recently alluded to in SARS-CoV-2 RNA-nucleoprotein-rich condensates^52^. Our model also accounts for the retention of the RdRP-bound viral transcripts inside the viral replicative condensates, in which the formation of inter-molecular RNA-RNA interactions between Seg1-Seg 11 transcripts is favoured in the presence of the viral RNA chaperone NSP2^6,26^.

During early infection, RNA-binding proteins NSP2 and NSP5 are the most abundant viral proteins produced in cells^12^, each being indispensable for the viral RNP condensate formation. The *K*_d_ value of the promiscuous RNA-binding protein NSP2 for ssRNA is low nanomolar^27,53^, thus protein-free RNAs would be expected to be bound by it, consistent with our previous observations of RV transcripts interacting with a non-viral RNA in the presence of NSP2 *in vitro*^26^. Thus, viral transcript sequestration within the specialized complex RNP coacervates offers a simple solution to the problem of co-assembling coding RV transcripts in the presence of other non-cognate RNAs, and ribosomes that may interfere with the RNA-RNA assembly process. Given the degenerative nature of RNA-RNA base-pairing, such non-specific interactions are likely to take place in cells^30^, and thus they would be expected to interfere with the stoichiometric assembly of eleven distinct viral transcripts. This implies that the unique viroplasmic condensate composition must be maintained through enrichment of cognate RNA-binding viral proteins. Remarkably, targeting the RV RNP condensates by a catalytically active Cas6f/Csy4 endonuclease fused to NSP5 results in the viral RNA genome editing that occurs within these condensates^22^, further confirming the essential role of these RNP condensates in the RV transcript selection and enrichment. The UDEx-FISH analysis of single cells revealed that the apparent stoichiometries of viroplasm-enriched and cytoplasmically-localised RV transcripts were different, suggesting that the observed RNA partitioning mechanism may also contribute to the efficient stoichiometric enrichment of eleven non-identical viral transcripts in these condensates. We propose that viroplasms may serve as the crucibles for the robust stoichiometric assembly of viral transcripts enriched in these RNA chaperone-rich condensates, which will underpin further studies to explore the exact mechanisms of RNA enrichment as a potential intervention strategy.

## METHODS

### Cells and viruses

MA104 (Clone 1, ATCC CRL-2378.1) cells (see **Supplementary Table S1**) were propagated and handled as previously described 54. MA104 cell line (*Cercopithecus aethiops kidney epithelial cells*) stably expressing NSP5-EGFP 18 was cultured in DMEM (Dulbecco’s modified Eagle medium, GlutaMax-I, 4.5 g/L glucose, ThermoFisher), supplemented with 10% foetal bovine serum (FBS), 1% non-essential amino acids solution (Sigma), 1 mM sodium pyruvate (Sigma) and 500 μg/ml G418 (Roche). Rotavirus A (RVA) strain RF (G6P6[1]) was a generous gift from Dr Ulrich Desselberger (University of Cambridge, UK). It was cultivated, harvested and stored, as described previously^54,55^. For RNA imaging experiments, MA104 cells and their derivatives (MA104-NSP5-EGFP), MA104-shRNA-NSP2) were seeded into Ibidi 8-well μ-slides and allowed to reach 90% confluency prior to the infection. Confluent cell monolayers were rinsed twice with DMEM medium without FBS for 15 minutes to remove any residual FBS, and were subsequently infected with trypsin-activated rotavirus stocks, as described in 54 at multiplicity of infection (MOI) of 10. Cells were fixed at different time points of infection, as described below.

### Generation of stable cell lines

MA104-shRNA-NSP2 cell line was generated using the PiggyBac system. Briefly, 10^5^ MA104 cells were co-transfected with the plasmid pCMV-HyPBase encoding the hyperactive variant of PiggyBac transposase^56,57,22^ along with the plasmid pPB[shRNA]-EGFP:T2A:Puro-U6 harbouring shRNA targeting RVA NSP2 gene using Lipofectamine 3000 (Sigma-Aldrich), following the manufacturer’s instructions. The cells were maintained in DMEM supplemented with 10% FBS for 3 days, and then the cells were subjected to selection in the presence of 5 μg/ml puromycin (Sigma-Aldrich) for 4 days, prior to further selection by FACS sorting for EGFP expression.

### Immunofluorescence (IF) detection of viral proteins

MA104 cells and their derivatives were grown on glass coverslips and infected with RVs, as described above. The cells were fixed with 4% (v/v) paraformaldehyde in PBS for 15 min at room temperature then rinsed twice with PBS and incubated for 10 min in a 0.1 M solution of glycine in PBS pH 7.4. Samples were then permeabilised with 0.2% Triton X-100 (PBS, v/v) for 10 min, rinsed with PBS and incubated in PBS containing 3% molecular biology grade BSA (New England Biolabs) and 0.05% Tween-20 for 1 h at room temperature. The cells were then incubated with guinea pig NSP2 antiserum (**Supplementary Table S1**) diluted 1:2000 in PBS containing 3% molecular biology grade BSA and 0.05% Tween-20, or 1:200 for a monoclonal anti-VP6 antibody (2B4, **Supplementary Table S1**) in PBS containing 3 % BSA and left overnight at +4°C. The cells were washed 3 times with PBS containing 0.1% Tween-20, followed by co-incubation with their respective anti-guinea pig and anti-mouse fluorescently labeled secondary antibodies (**Supplementary Table S1**), following the manufacturer’s instructions. Cells were washed 3 times with PBS and coverslips were mounted with Vectashield Antifade Reagent containing DAPI (ThermoFisher).

### Western blot analysis

Proteins were treated with SDS and 2-mercaptoethanol at 95°C, resolved by SDS-PAGE on Tris-glycine gels and transferred onto nitrocellulose membranes. Afterwards, the membranes were blocked with PBS containing 5% (w/v) skimmed milk and 0.1 % Tween-20 and then incubated with guinea pig NSP5-specific antibody (**Supplementary Table S1**) dilluted in PBS containing 1% milk and 0.1% Tween-20. Blots were washed with PBS and incubated with HRP-conjugated secondary antibodies in PBS containing 1 % milk and 0.1 % Tween-20. Blots were developed with SuperSignal West Pico Chemiluminescent Substrate (Pierce) and exposed to BioMax MR film (Kodak). All scanned images were post-processed in Adobe Illustrator.

### Recombinant NSP3-2A-EGFP virus

Rescue of the recombinant NSP3-2A-EGFP virus (strain SA11) was carried out as previously described^22,31,58,59^. Briefly, monolayers of BHK-T7 cells (4 × 10^5^) cultured in 12-well plates were co-transfected using 2.5 μL of TransIT-LT1 transfection reagent (Mirus) per microgram of DNA plasmid. Each mixture comprised 0.8 μg of SA11 rescue plasmids: pT_7_-VP1, pT_7_-VP2, pT_7_-VP3, pT_7_-VP4, pT_7_-VP6, pT_7_-VP7, pT_7_-NSP1, pT_7_-NSP3-2A-EGFP, pT_7_-NSP4, and 2.4 μg of pT_7_-NSP2 and pT_7_-NSP5. 0.8 μg of pcDNA3-NSP2 and 0.8 μg of pcDNA3-NSP5, encoding NSP2 and NSP5 proteins, were also co-transfected to increase the efficiency of virus rescue. At 24 h post-transfection, MA104 cells (5 × 10^4^ cells) were added to transfected cells. The cells were co-cultured for 3 days in FBS-free medium supplemented with porcine trypsin (0.5 μg/mL) (Sigma Aldrich). After incubation, transfected cells were lysed by freeze-thawing and 0.2 ml of the lysate was used to infect fresh MA104 cells. After adsorption at 37°C for 1 hour, cells were washed three times with PBS and further cultured at 37°C for 4 days in FBS-free DMEM supplemented with 0.5 μg/mL trypsin (Sigma Aldrich, 9002-07-7) until a clear cytopathic effect was visible. Successful production of EGFP by the recombinant virus was confirmed microscopically.

### Single-molecule Fluorescence in situ Hybridisation (smFISH)

Rotavirus-infected and mock-infected MA104 cell controls, where appropriate, were fixed with 4% (v/v) methanol-free paraformaldehyde in nuclease-free phosphate saline buffer (PBS) for 10 min at room temperature. Samples were then washed twice with PBS, and fixed cells were permeabilised with 70% (v/v) ethanol (200 proof) in RNAse-free water, and stored in ethanol at +4°C for at least 12 hours prior to hybridization, and no longer than 24 h. Permeabilized cells were then re-hydrated for 5 min in a pre-hybridization buffer (300 mM NaCl, 30 mM trisodium citrate, pH 7.0 in nuclease-free water, 10 % v/v Hi-Di formamide (Thermo Scientific), supplemented with 2 mM vanadyl ribonucleoside complex). Re-hydrated samples were hybridized with an equimolar mixture of DNA probes specific to the mRNA targets (RVA RF or *C.aethiops* GAPDH transcripts), 62.5 nM final concentration, see Supplementary file 1, in a total volume of 200 μl of the hybridization buffer (Stellaris RNA FISH hybridization buffer, Biosearch Technologies, supplemented with 10% v/v Hi-Di formamide). After 4 hours of incubation at 37°C in a humidified chamber, samples were briefly rinsed with the wash buffer (300 mM NaCl, 30 mM trisodium citrate, pH 7.0, 10 % v/v formamide in nuclease-free water, after which a fresh aliquot of 300 μl of the wash buffer was applied to each well and incubated twice at 37°C for 30 min. After washes, nuclei were briefly stained with 300 nM 4’,6-diamidino-2-phenylindole (DAPI) solution in 300 mM NaCl, 30 mM trisodium citrate, pH 7.0) and the samples were finally rinsed with and stored in the same buffer without DAPI prior to the addition of photostabilising imaging buffer (PBS containing an oxygen scavenging system of 2.5 mM protocatechuic acid, 10 nM protocatechuate-3,4-dioxygenase supplemented with 1 mM (±)-6-hydroxy-2,5,7,8-tetramethylchromane-2-carboxylic acid (Trolox) 60.

Sequences of the oligonucleotide RNA FISH probes, as listed in Supplementary file 1 were used. These were generated using the Stellaris RNA FISH probe designer (https://www.biosearchtech.com/stellaris-designer), using each gene-specific ORF sequences (see Supplementary file 1 for GenBank IDs) and level 2 masking. The resulting pools of probes were then further filtered to remove the sequences targeting the RNA transcripts sequences with higher propensity to form stable intra-molecular base-pairing.

### Universal DNA Exchange FISH (UDEx-FISH)

For UDEx-FISH, sequences of the oligonucleotide RNA FISH probes are listed in Supplementary file 1. Gene-specific portions of each probe were generated using the Stellaris RNA FISH probe designer, followed by a TT linker and a 10-nt long DNA handle designed to minimize any potential intra-molecular base-pairing 23. All FISH hybridization steps were identical to the ones described in the section above, except the individual target-specific probe concentration was adjusted to 62.5 nM in the hybridization mix. Cy3- and Cy5-modified labeling DNA strands (‘Imagers’) were incubated in the labeling buffer (600 mM NaCl, 2.7 mM KCl, 8 mM Na_2_HPO_4_ and 2 mM KH_2_PO_4_ pH 7.4 in nuclease-free water), for approximately 5 min. Then, samples were rinsed with the labeling buffer, followed by addition of the imaging buffer (*vide supra*). After a round of image acquisition, hybridized labeling strands were dissociated by briefly incubating samples in 30% (v/v) formamide in PBS (‘Wash buffer’) for 2-3 min at RT, repeating this procedure twice, whilst monitoring for any residual fluorescence signal in four acquisition channels (Cy5, Cy3, GFP, DAPI). Formamide-containing wash buffer was then aspirated and samples were rinsed twice with fresh labeling buffer prior to the introduction of the next batch of ‘Imager’ DNA strands, thus concluding a single imaging cycle. To calibrate the recorded fluorescence intensities to quantify the signals originating from the individual transcripts in an unbiased manner due to differences in properties of spectrally distinct dyes (Cy3 vs Cy5), we swapped Cy5 and Cy3-dye labelled DNA imagers for one of the RNA targets (**Supplementary Fig. S7, b**) in each imaging cycle. A total number of 6 imaging cycles were required to image the RV transcriptome, and the last cycle always contained an Imager targeting the segment imaged during cycle 1 to control for a possibility of the signal loss due to the dissociation of FISH probes, and for signal calibration purposes.

### Image data acquisition

Widefield imaging was carried out on a Leica (Wetzlar, Germany) DMI6000B inverted microscope equipped with a LEICA HCX PL APO 63x/NA1.4 oil immersion objective. Dye excitation was performed with a cooled pe-4000 LED source illumination system at the wavelengths 385 nm (DAPI), 470 nm (eGFP), 550 nm (Cy3 and Quasar 570) and 635 nm (Cy5 and Quasar 670).

Fluorescent signals were detected with a Leica DFC9000 GT sCMOS camera with a pixel size of 6.5 μm. Images were acquired over a full field of view of the camera chip (2048 × 2048 pixels) resulting in a total imaging region of 211 μm × 211 μm. Exposure times were adjusted accordingly to the signal intensity to avoid pixel saturation. Typical exposure times were 250 ms for DAPI, 500 ms for GFP, 750 ms – 1 s for Cy3/Cy5 or Quasar 570/670 dyes. Image stacks were acquired using 250 nm Z-axis steps across a range of approximately 5 μm. Full stacks were recorded consecutively for each channel, from the lowest to the highest energy excitation wavelength.

Confocal imaging was carried out on a Zeiss (Jena, Germany) LSM780 confocal laser-scanning microscope equipped with a Zeiss Plan-APO 63×/1.46-NA) oil-immersion objective. DAPI, GFP, Cy3B excitation were performed with 405 nm, 488 nm and 561 nm lasers, respectively, and the pinhole sizes were adjusted to 1 Airy unit. Samples were immersed in Vectashield (H-1000, Vector Laboratories) prior to imaging. Image stacks were acquired using 377 nm Z-axis steps across a range of approximately 5 μm. Image acquisition was controlled using the Zeiss Zen Black Software (Zeiss).

DNA-PAINT imaging was carried out on an inverted Nikon Eclipse Ti microscope (Nikon Instruments) equipped with the Perfect Focus System using objective-type total internal reflection fluorescence (TIRF) configuration (oil-immersion Apo SR TIRF, NA 1.49 100x objective). A 200 mW 561 nm laser beam (Coherent Sapphire) was passed through a cleanup filter (ZET561/10, Chroma Technology) and coupled into the microscope objective using a beam splitter (ZT561rdc, Chroma Technology). Fluorescence light was spectrally filtered with an emission filter (ET575lp, Chroma Technology) and imaged with an sCMOS camera (Andor Zyla 4.2) without further magnification, resulting in an effective pixel size of 130 nm after 2×2 binning. Images were acquired using a region of interest of 512×512 pixels. The camera read-out rate was set to 540 MHz and images were acquired with an integration time of 200 ms. 5’-ATACATTGA-Cy3B-3’ was used as imager strand sequence. Further details of imaging conditions for each experiment are presented in **Supplementary Table S2**.

### Image Processing and Colocalisation Analysis

Deconvolution analysis was applied to all acquired widefield images using the Huygens Essential software (Scientific Volume Imaging B.V., the Netherlands). All channels and Z-planes were deconvolved using Huygens batch express tool (Standard profile).

Z-stacks were loaded with ImageJ and out-of-focus planes were manually discarded. Prior to analysis a maximum intensity Z-projection was performed with the remaining Z-planes.

2D colocalization analysis was performed using maximum intensity Z-projections analysed in Icy (Version 1.9.10.0), an open bioimage informatics platform (http://icy.bioimageanalysis.org/). Regions of interest were drawn around individual cells prior to the analysis. Pearson’s correlation coefficient (PCC) were chosen as a statistic for quantifying colocalization to measure the pixel-by-pixel covariance in the signal levels between two distinct channels. This allows subtraction of the mean intensity from each pixel’s intensity value independently of signal levels and the signal offset for each ROI. Pearson’s correlation coefficient values were calculated in Icy Colocalization Studio that employs pixel scrambling method^61^.

### DNA-PAINT image analysis

Raw fluorescence data were subjected to super-resolution reconstruction using Picasso software package^24,39^. Drift correction was performed with a redundant cross-correlation and gold particles used as fiducial markers. The apparent on-rate (*apparent k*_*on*_) of imager stands binding to their corresponding docking sites was used to quantify the relative number of binding sites. A higher *apparent k*_*on*_ value indicates a higher number of binding sites, i.e., RNA molecules detected in the structure^25^. *k*_*on*_ values were calculated for each selected structure assuming *k*_*on*_ = (*τ*_*D*_ × *c*)^−1^, where *c* is the imager strand concentration and *τ*_*D*_ the dark time between binding events^25^. The number of FISH probes per RNA was calculated assuming *k*_*on*_ = 1 × 10^6^(*Ms*)^−1^ for each docking site. Further quantification and fitting were performed using OriginPro, as previously described^25^.

### Single-Cell RNA Imaging and Viral Transcriptome Analysis using UDEx-FISH

Images were recorded with the same acquisition parameters for all rounds. Integrated signal densities and areas of single cells were measured in ImageJ to calculate signal intensities and areas for each RNA target. Background signals were determined from signals measured in mock-infected cells, and these were subtracted for each channel respectively. Signals in Cy3 and Cy5 channels were calibrated by calculating the correction factors for Cy3/Cy5 signals for one of the RNA targets that was imaged sequentially in both channels using Cy3 and Cy5 imager strands.

### Host Cell and Viral Transcriptome RNA-Seq Data Analysis

RV-infected MA104 cells were harvested at 6 HPI, and total RNA was extracted using RNEasy kit (QIAGEN). 1 μg of total RNA was depleted of rRNA using NEBNext rRNA Depletion Kit (Human/Mouse/Rat) prior to cDNA synthesis primed with random hexanucleotide oligonucleotides (Random Primer 6, NEB). Sequencing library construction was carried out using NEBNext Ultra II FS DNA Library Prep Kit for Illumina (NEB), and the resulting library was sequenced using Illumina MiSeq v2 (2×150 bp) platform. Paired-end run raw sequencing data were pre-processed with fastp (v0.20.1) using default command line parameters^62^ to yield 1,625,571 reads. A bwa-mem2 (v2.1) index was generated (default parameters) using bovine rotavirus A strain RF reference genome (J04346.1, KF729639.1, KF729643.1, KF729690.1, KF729653.1, K02254.1, KF729659.1, Z21640.1). Identified viral transcript reads had 99.81% identity against the reference genome. These preprocessed reads were mapped to the index using bwa-mem2 mem^63^. All mapped reads were sorted using samtools sort (v1.11) to create a consensus structure of all reads using bcftools (v1.11)64,65. bcftools mpileup (command line parameters: -Ob -d 10000) was used to generate genotype likelihoods, followed by bcftools call (command line parameters: -Ob -mv) for SNP and indel calling, bcftools norm (command line parameters: -Ob) for indel normalization, bcftools index for indexing, and bcftools consensus was used to create the consensus structure. The consensus rotavirus RF genome file was combined with the transcriptome data available for *Chlorocebus sabaeus* sp. (MA104 host cell line) from Ensemble release 103 66 to construct a combined transcriptome. This combined transcriptome file was used to generate a salmon (v1.4.0) index (command line parameters: --keepDuplicates) to quantify the pre-processed Illumina reads using salmon quant [6] (command line parameters: -l A --validateMappings)67. A total of 671,700 reads were identified by salmon (i.e., estimate of the number of reads mapping to each transcript that was quantified), out of which 114,276 reads were mapped to the viral transcriptome. SRA Illumina reads data are available under the accession number PRJNA702157 (SRR13723918, RNA-Seq of Bovine Rotavirus A: Strain RF).

## Acknowledgements

We are indebted to Ulrich Desselberger (University of Cambridge, UK), Elena Conti (Max Planck Institute of Biochemistry, Munich) and Jack Bravo for their valuable comments and suggestions during the preparation of the manuscript. We would like to thank Ulrich Desselberger (University of Cambridge, UK) and Oscar R. Burrone (ICGEB, Trieste, Italy) for the generous gifts of the G6P6[1] strain RF of group A rotavirus, and for a recombinant NSP3-2A-EGFP rotavirus, respectively. The authors would like to thank Martin Spitaler (MPIB, Munich) for his excellent technical assistance with imaging samples. This work was supported by the Wellcome Trust (grants 103068/Z/13/Z and 213437/Z/18/Z to A.B.), ERC through an ERC Starting Grant (MolMap, Grant agreement number 680241), and the DGF through the SFB1032 (Nanoagents for the spatiotemporal control of molecular and cellular reactions, Project A11), the Max Planck Society, the Max Planck Foundation, and the Center for Nanoscience (CeNS). S.S. acknowledges support from the DFG through the Graduate School of Quantitative Biosciences Munich (QBM). This research was funded in part by the Wellcome Trust [213437/Z/18/Z]. For the purpose of Open Access, the author has applied a CC BY public copyright license to any Author Accepted Manuscript version arising from this submission.

## Supplementary Information

**Supplementary Figure S1.**
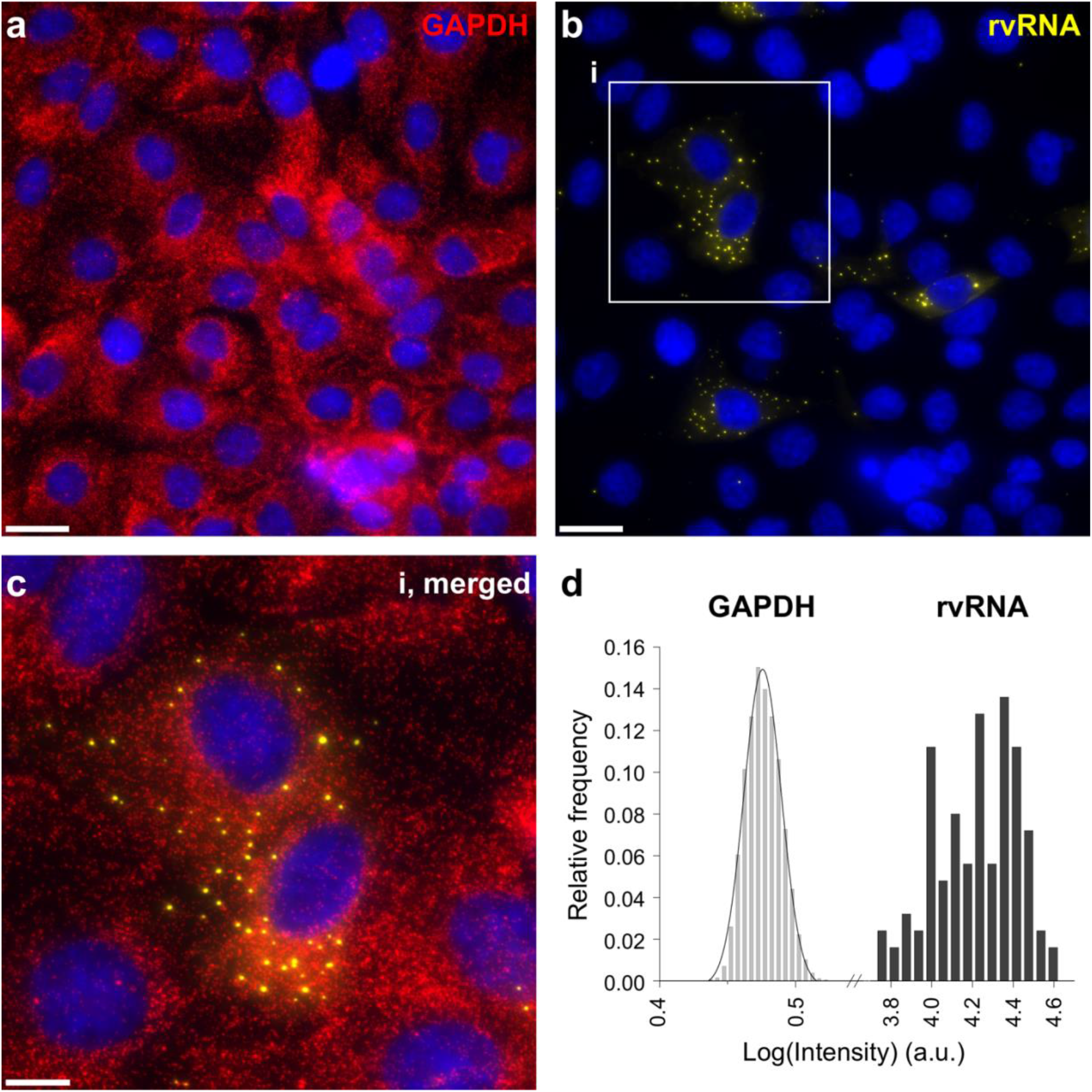
smFISH of GAPDH mRNA and RV transcripts. GAPDH mRNA (red, **a**) and RV transcripts (yellow, **b**) imaged using two distinct sets of FISH probes targeting GAPDH mRNA (Quasar 670 dye-labelled probes) and RV transcripts (Quasar 570 dye-labelled probes), as shown in Figure 1a. At 4 hpi, RV transcripts form distinct, high intensity foci. Scale bars, 25 μm. (**c**) A close-up view of the highlighted area (**i**). Scale bar, 10 μm. (**d**) Signal intensity distribution histograms for GAPDH transcripts and RV transcript foci (rvRNA).

**Supplementary Figure S2.**
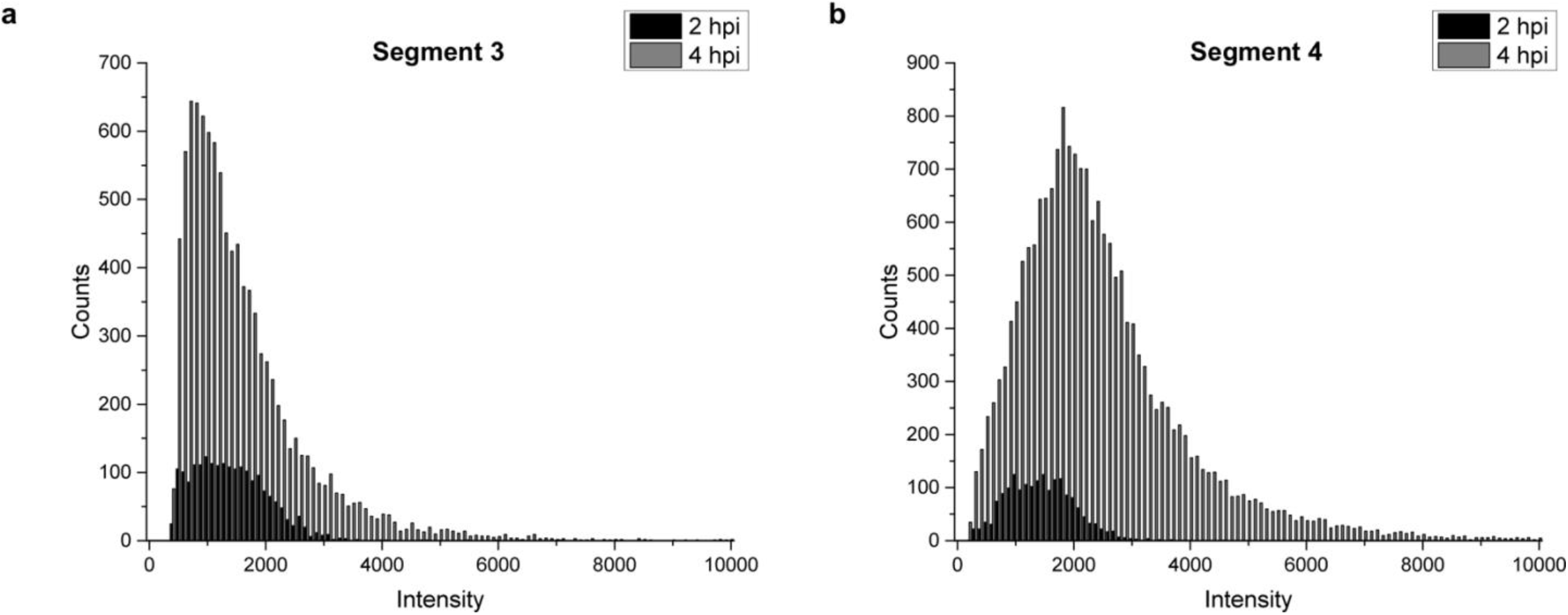
Signal intensity distributions of Seg3 and Seg4 transcript foci at 2 and 4 hpi. Raw signal intensity distributions (counts) of diffraction-limited smFISH-detected spots were derived for Seg3 (Quasar 670 dye-labelled probes) and Seg4 (Quasar 570 dye-labelled probes) transcripts at 2 and 4 hpi. Each spot was detected after applying a constrained deconvolution algorithm (see Materials and Methods). Uneven Gaussian widefield illumination patterns, and differences in excitation and detection efficiencies at variable z-planes, as well as different hybridization efficiencies of smFISH probes may also contribute to the observed broadening of the intensity distributions.

**Supplementary Figure S3.**
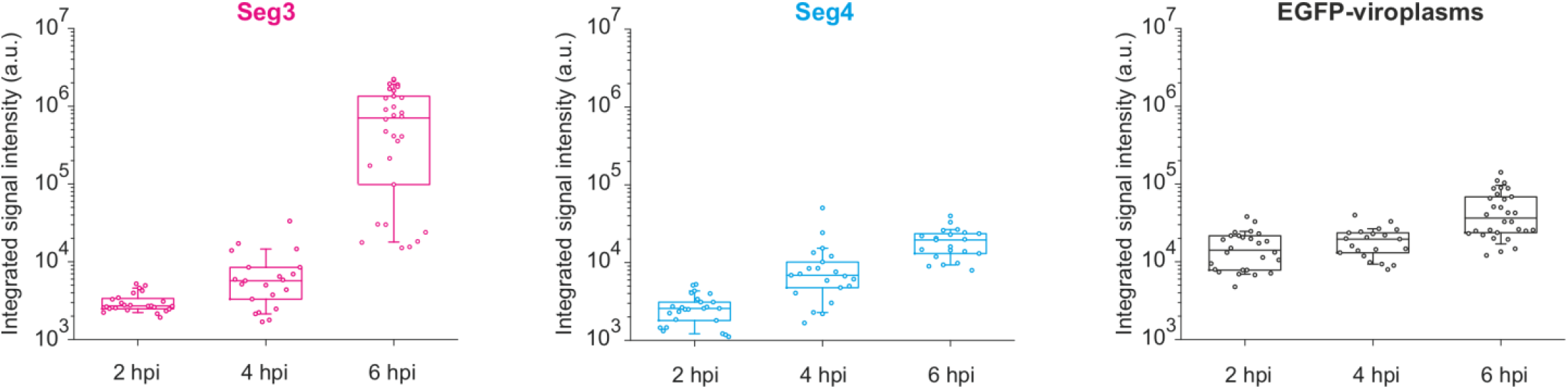
Integrated signal intensities of Seg3, Seg4 and EGFP-NSP5 at different RV infection time points. Changes in the integrated signal intensities for Seg3 (magenta), Seg4 (cyan) RNAs and EGFP-tagged viroplasms (black) over the course of RV infection. Integrated signals were calculated for single cells separately, N = 37 (2 hpi), N = 62 (4 hpi), N = 51 (6 hpi).

**Supplementary Figure S4.**
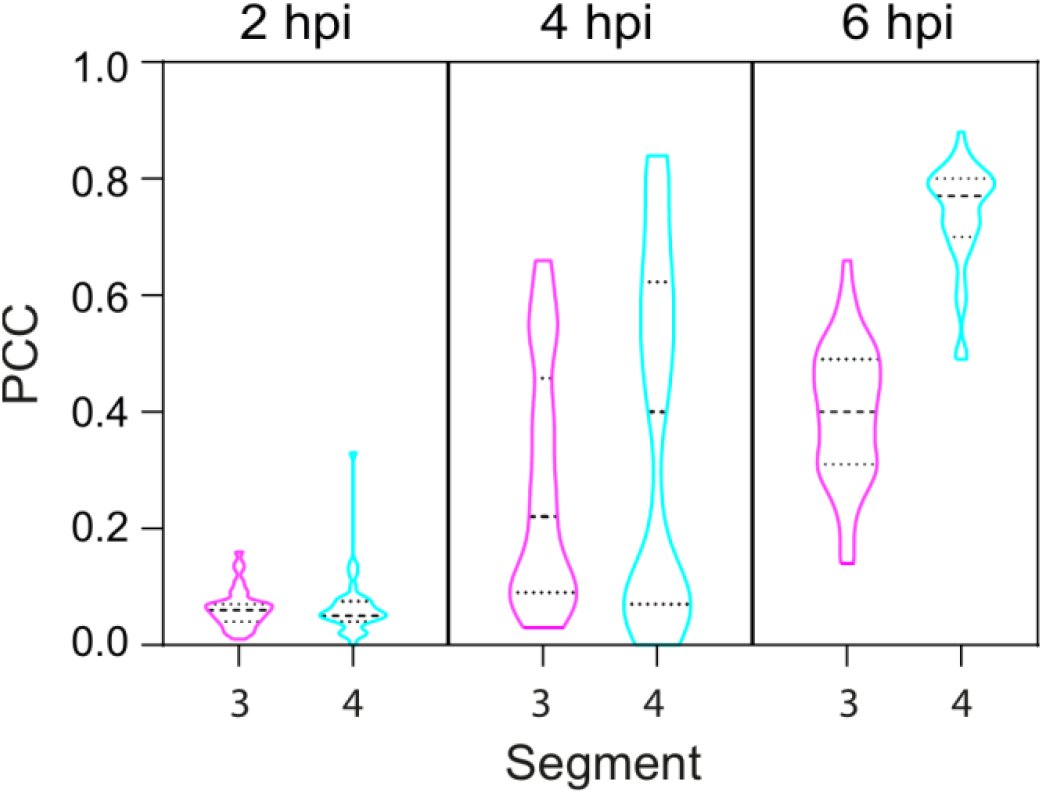
Colocalisation analysis of EGFP-NSP5 with Seg3 and Seg4. Colocalisation of EGFP-NSP5-tagged viral factories with Seg3 (magenta violin plots) and Seg4 (cyan violin plots) transcripts over the course of RV infection. Median and quartile values of the calculated Pearson’s correlation coefficients (PCCs) are shown as dashed and dotted lines, respectively. Each data point represents the PCC value calculated for a single cell. N = 37 (2 hpi), N = 62 (4 hpi), N = 51 (6 hpi).

**Supplementary Figure S5.**
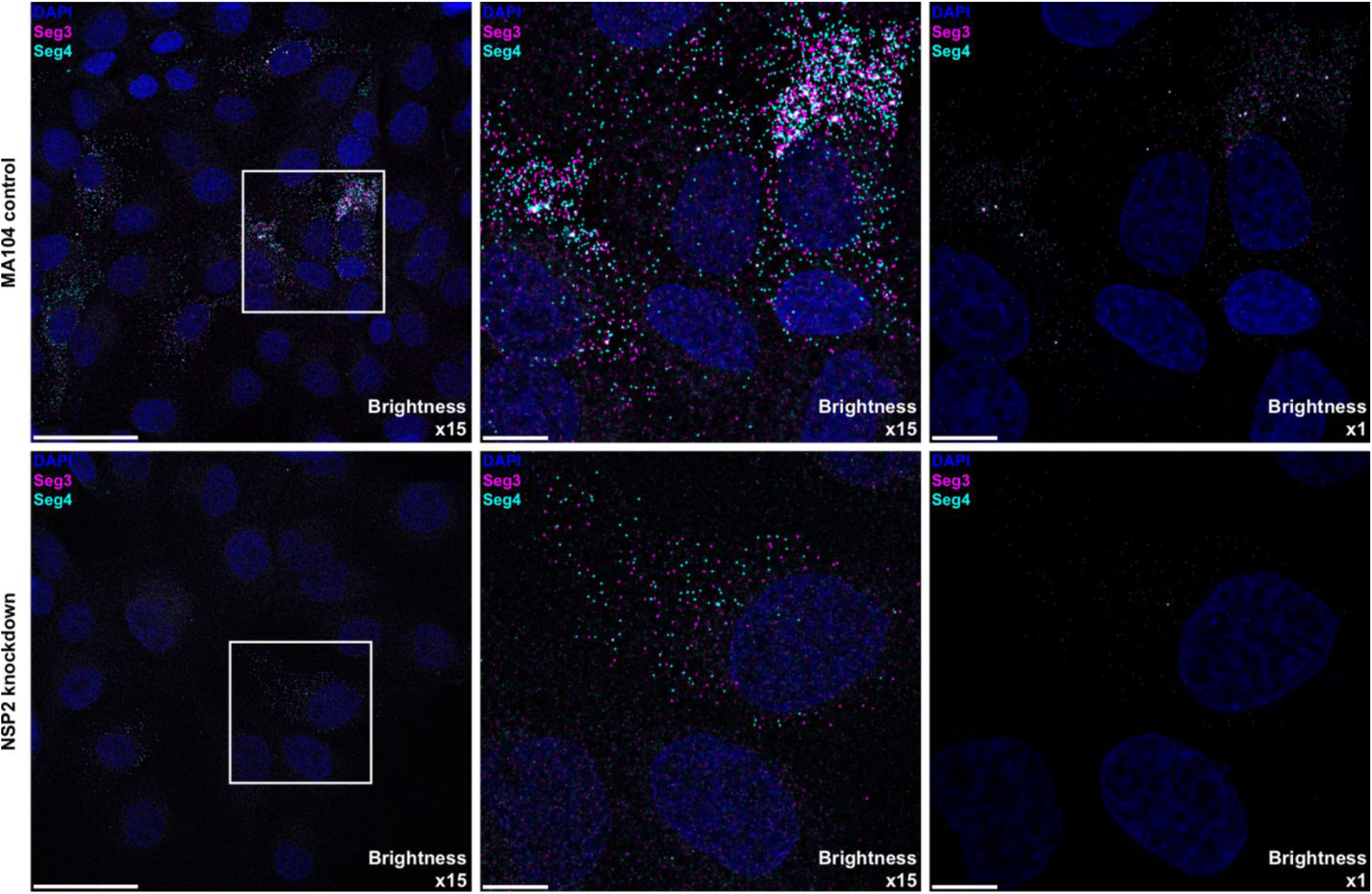
NSP2 knockdown in MA104 cells results in the apparent loss of RNA oligomerisation. Note that in both cases single Seg3 and Seg4 transcripts accumulate in the cytosol, while the signal intensity of RNAs in case of NSP2 knockdown does not increase over time. Scale bars, 50 μm (left), 10 μm (middle & right panel).

**Supplementary Figure S6.**
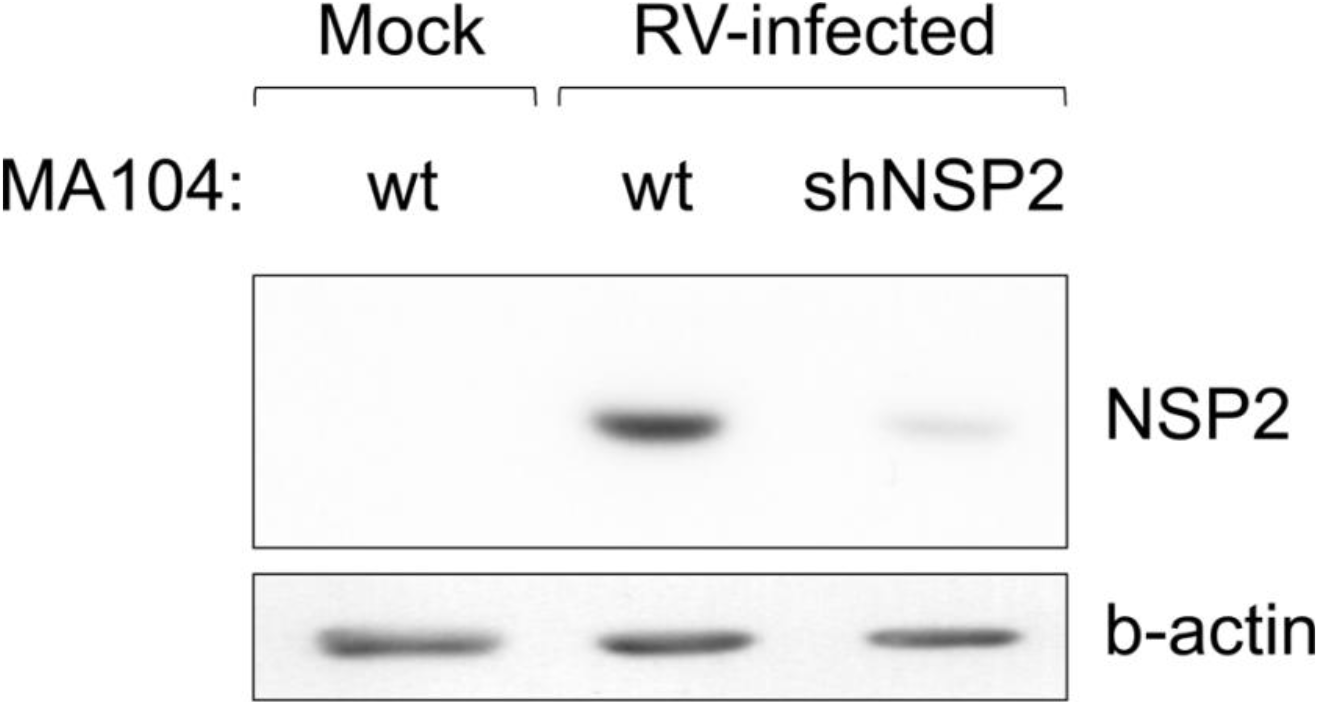
Western blot of NSP2 shRNA knock down. shRNA knock-down of NSP2 in RV-infected MA104 cells analysed by smFISH shown in the Supplementary Figure 6. MA104 cells stably expressing shRNA targetting NSP2 transcripts were infected with MOI of 5 and cells were analysed for NSP2 expression by Western blot.

**Supplementary Figure S7.**
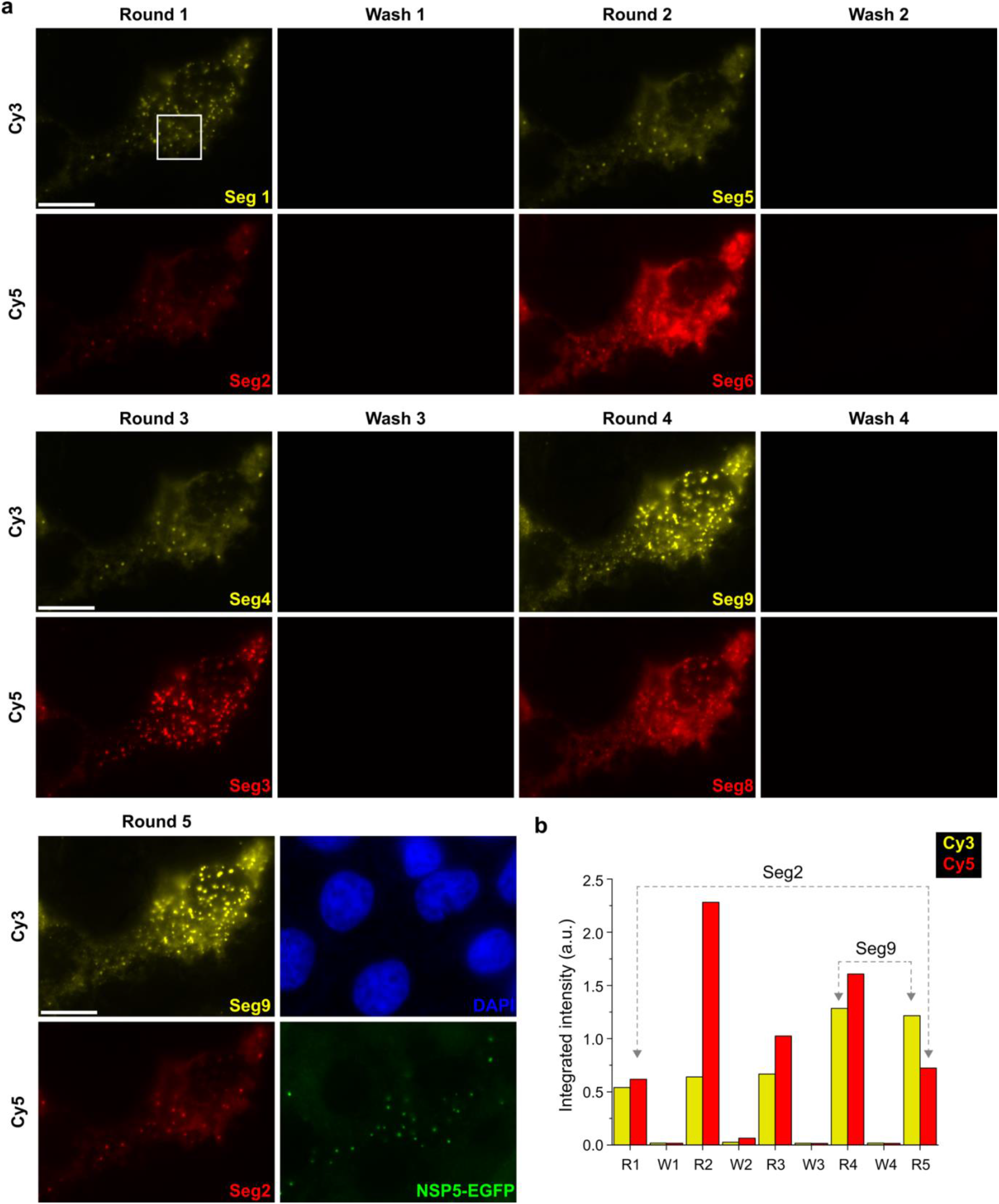
Evaluation of the UDEx-FISH approach in RV-infected MA104 cells. (**a**) Washing and labeling efficiencies of smFISH probes with orthogonal DNA handles designed for Segments 1-9. After each wash step no residual RNA signal was detected for each segment tested in this experiment. In addition, Seg2 target was then re-imaged iteratively to ensure that multiple washing steps did not dissociate FISH probes pre-hybridized to a target RNA. (**b**) The integrated signal density of the after each successive round of imager dissociation and re-association. Each image plane for each channel was recorded with the same image acquisition parameters. Brightness levels were set identical for each color channel for all the images shown. Scale bars, 15 μm.

**Supplementary Table S1.**
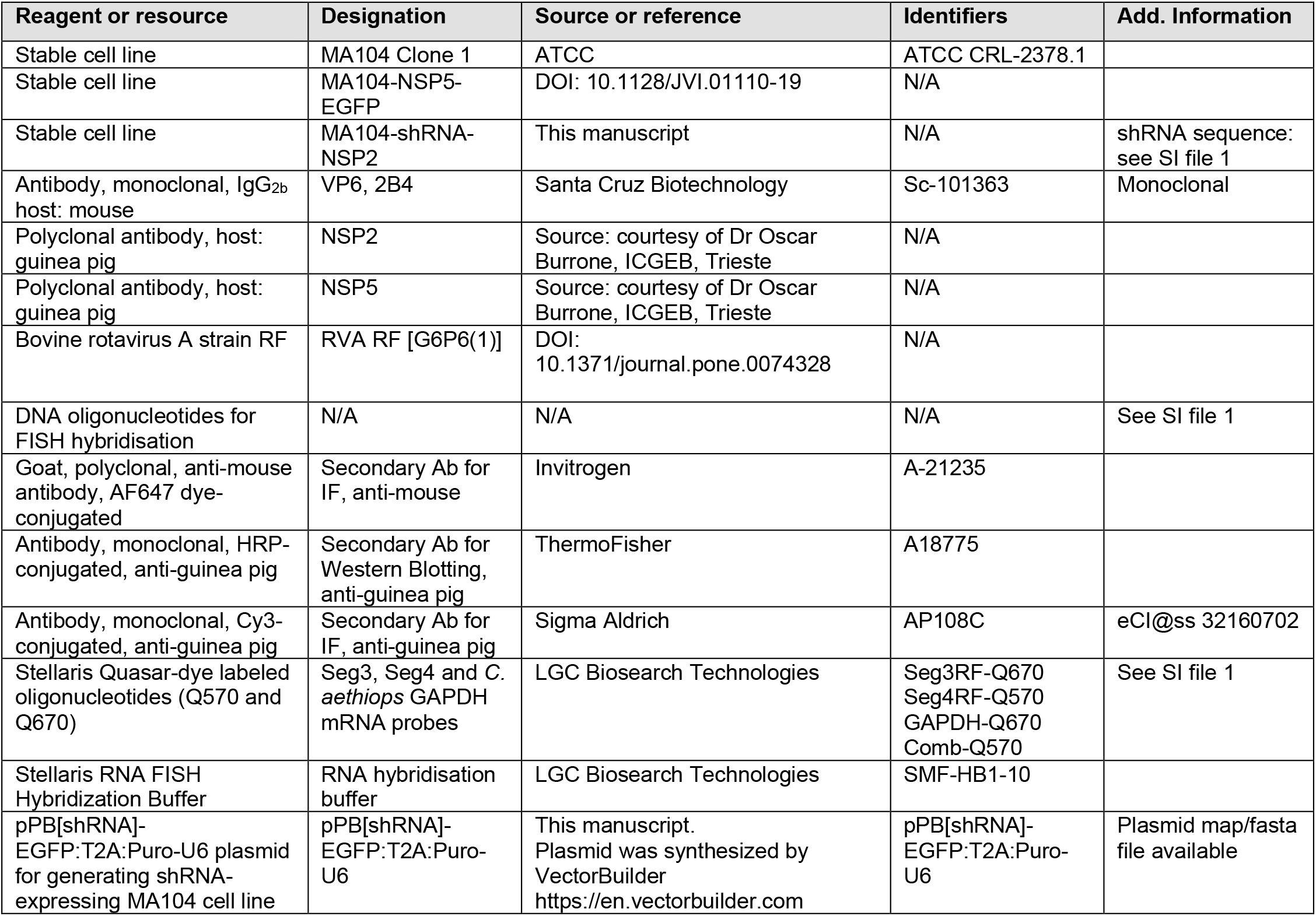
Key resource table.

**Supplementary Table S2.**
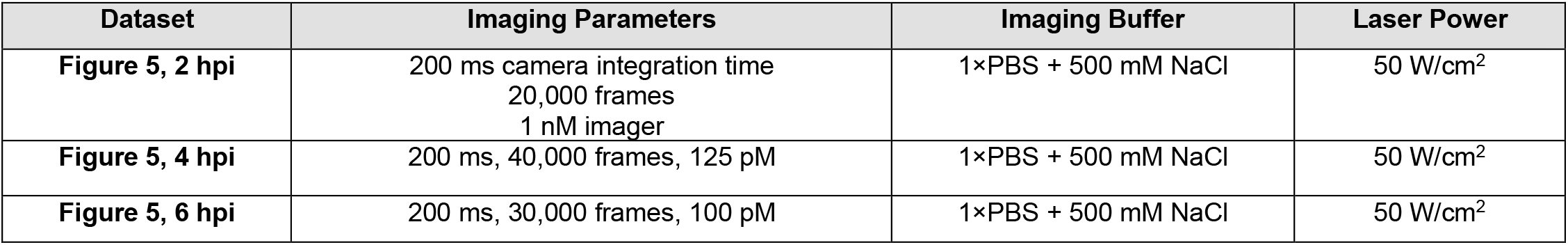
DNA-PAINT imaging conditions.

